# SARS-CoV-2 proteins and anti-COVID-19 drugs induce lytic reactivation of an oncogenic virus

**DOI:** 10.1101/2020.10.02.324228

**Authors:** Jungang Chen, Lu Dai, Lindsey Barrett, Steven R. Post, Zhiqiang Qin

## Abstract

An outbreak of the novel coronavirus SARS-CoV-2, the causative agent of Coronavirus Disease-2019 (COVID-19), a respiratory disease, has infected over 34,000,000 people since the end of 2019, killed over 1,000,000, and caused worldwide social and economic disruption. Due to the mechanisms of SARS-CoV-2 infection to host cells and its pathogenesis remain largely unclear, there are currently no antiviral drugs with proven efficacy nor are there vaccines for its prevention. Besides severe respiratory and systematic symptoms, several comorbidities may also increase risk of fatal disease outcome. Therefore, it is required to investigate the impacts of COVID-19 on pre-existing diseases of patients, such as cancer and other infectious diseases. In the current study, we have reported that SARS-CoV-2 encoded proteins and some anti-COVID-19 drugs currently used are able to induce lytic reactivation of Kaposi’s sarcoma-associated herpesvirus (KSHV), one of major human oncogenic viruses through manipulation of intracellular signaling pathways. Our data indicate that those KSHV+ patients especially in endemic areas exposure to COVID-19 or undergoing the treatment may have increased risks to develop virus-associated cancers, even after they have fully recovered from COVID-19.

## Introduction

COVID-19 has become a devastating pandemic since its origin in the city of Wuhan, Hubei province of China in December 2019 (Zhu et al., 2020; Wu et al., 2020). Based on the updated information from Johns Hopkins Coronavirus Resource Center, there are now more than 34 million global confirmed cases and over 1 million deaths. In the United States, the confirmed cases and deaths have reached 7.2 million and 207,000, respectively. In addition, the COVID-19 pandemic has caused a huge economic loss due to the necessary shut down and quarantine procedures. Further, many countries have seen secondary spikes in case numbers after reopening their economies. Septic shock and multiple organ failure represent the most common immediate causes of death in patients with severe COVID-19. These deaths are mostly due to suppurative pulmonary infection, onset of cytokine storms, and the direct attack on multiple organs (Elezkurtaj et al., 2020). However, several comorbidities, such as hypertension, cardiovascular disease, endocrine disorder, diabetes and obesity may increase the likelihood of death from COVID-19 infection (Bennett et al., 2020). Therefore, it is very meaningful to investigate the impacts of COVID-19, including its relationship to the pre-existing diseases of patients, such as cancer and other infectious diseases.

KSHV is the etiologic agent of several human cancers including Kaposi’s sarcoma (KS), Primary effusion lymphoma (PEL), and Multicentric Castleman’s disease (MCD) (Broussard and Damania, 2020a), which are mostly seen in immunosuppressed patients. KS is an endothelial-originated multicentric malignant neoplasm, while PEL and MCD represent two kinds of B-cell lineage disorders (Polizzotto et al., 2017). This oncogenic virus belongs to the human γ-herpesvirus subfamily, and has two alternating life-cycle programs following primary infection in host cells, the latent and lytic phases (Broussard and Damania, 2020b). During latency, viral genomes persist as circular episomes with no progeny virion produced and only a limited number of latency-associated genes are expressed. Once entering the lytic phase, almost all viral genes are expressed, followed by the replication and release of mature virions. Recent findings have indicated that both viral latent and lytic proteins can play a pivotal role in the initiation and progression of virus-associated cancers (Aneja and Yuan, 2017). Here we try to understand if COVID-19 infection and the related treatments affect KSHV replication and increase the risk of developing virus-associated cancers, and how these mechanisms work.

## Results and Discussion

The human iSLK.219 cell line carries a recombinant rKSHV.219 virus encoding a constitutive GFP reporter and a RTA-inducible RFP reporter in the viral genome, thereby facilitating the monitoring of viral maintenance and lytic reactivation (Myoung and Ganem, 2011). To test potential impacts on KSHV replication by SARS-CoV-2, the iSLK.219 cells were transfected with a vector control or vectors encoding two of SARS-CoV-2 major structural proteins, spike protein (S) and nucleocapsid protein (N), and KSHV-RTA (a key viral protein controlling the latency to lytic switch, used as a positive control), respectively, with or without a low dose of doxycycline (Dox at 0.1 μg/mL) for 72 h induction. The ectopic expression of SARS-CoV-2 S and N proteins have been confirmed by immunoblots (**Fig. S1**). Under the condition without Dox induction, transfection of KSHV-RTA induced only a few cells to undergo lytic reactivation. In contrast, with low dose of Dox induction, transfection of either SARS-CoV-2 S or N vectors greatly increased lytic reactivation when compared to vector control, although a little less than KSHV-RTA (**Fig. 1A**). To confirm these results, we also transfected the same vectors into BCP-1, a KSHV+ PEL cell line. Our qRT-PCR data showed that transfection of either SARS-CoV-2 S or N vectors significantly increased representative lytic gene expression (Immediate-early gene, RTA; Early gene, ORF59; and Late gene, ORF17) when compared to vector control, regardless of with or without TPA induction (**Fig. 1B**). Considered together, these data indicate that SARS-CoV-2 has the potential to induce KSHV lytic reactivation.

**Figure 1.**
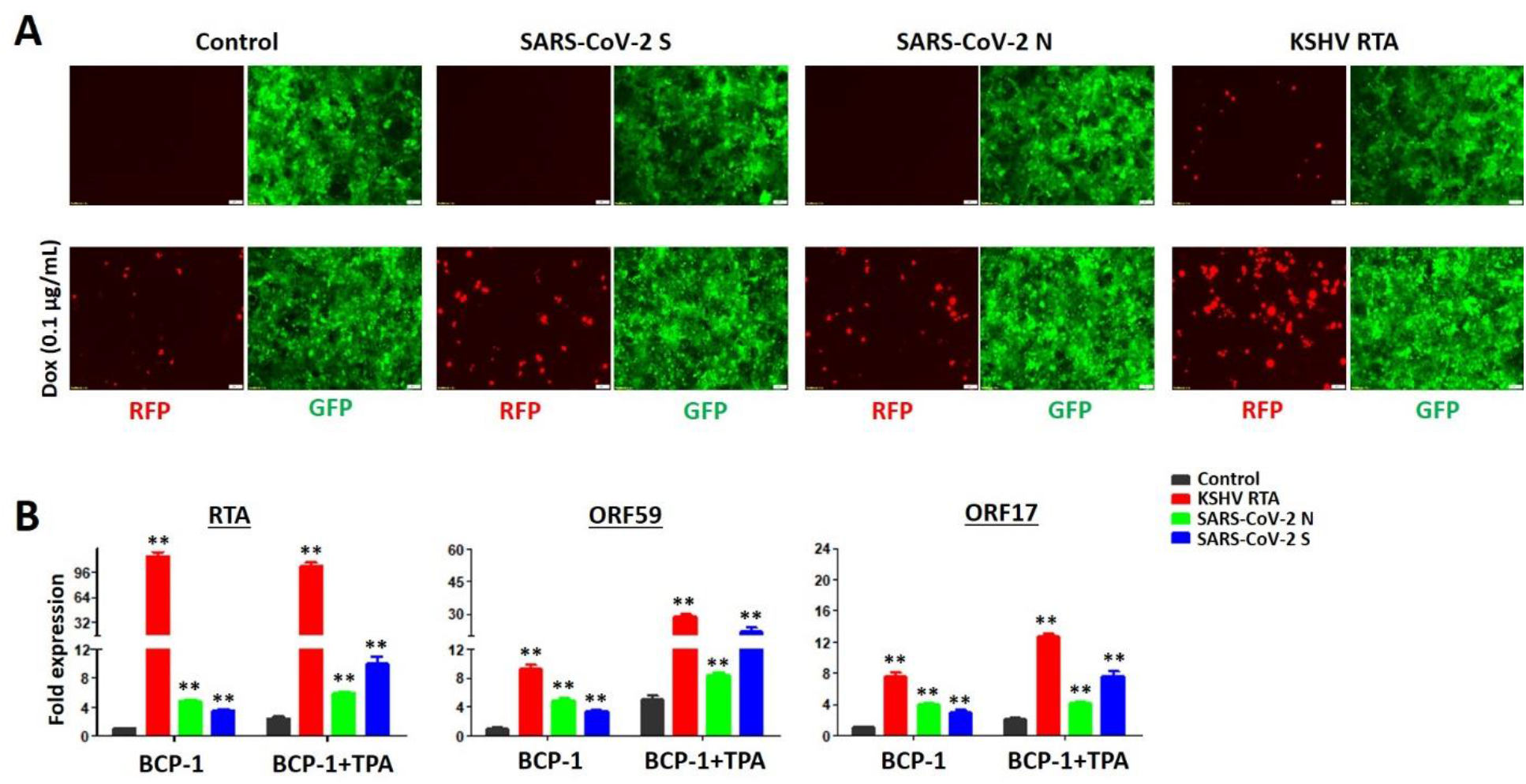
Ectopic expression of SARS-CoV-2 proteins induces KSHV lytic gene expression from latently infected cells. (**A**) The iSLK.219 cells were transfected with vector control or vectors encoding SARS-CoV-2 spike protein (S), nucleocapsid protein (N) and KSHV RTA (as a positive control) with or without low dose of doxycycline (Dox, 0.1 μg/mL) induction for 72 h. The expression of RFP (representing viral lytic reactivation) and GFP (representing infected cells) were detected using fluorescence microscopy. (**B**) BCP-1 cells were transfected as above with or without low dose of 12-*O*-tetradecanoyl-phorbol-13-acetate (TPA, 1.0 ng/mL) induction for 72 h, then the transcripts of representative lytic genes were quantified by using qRT-PCR. Error bars represent S.D. for 3 independent experiments, ** = p<0.01 (*vs* the vector control).

We next examined whether some drugs currently used for treatment of COVID-19 patients may affect KSHV lytic reactivation. A total 6 drugs reported were used in our study, including Azithromycin, Chloroquine diphosphate, Hydroxychloroquine sulfate, Nafamostat mesylate, Remdesivir, and Tocilizumab (Fiolet et al., 2020; Osawa et al., 2020; Maciorowski et al., 2020; Schoot et al., 2020). Through screening with iSLK.219 cells, we identified that two of these drugs, Azithromycin (an antibiotic) and Nafamostat mesylate (a synthetic serine protease inhibitor), greatly induced KSHV lytic reactivation in a dose-dependent manner under the condition with low dose Dox induction (**Figs. 2A & S2**). Furthermore, both Azithromycin and Nafamostat mesylate treatment promoted the production of mature virions as detected using an infectivity assay in which the supernatants from induced iSLK.219 cells were used to infect HEK293T cells. As shown in **Fig. 2B**, the GFP signal in HEK293T cells demonstrates successful production of infectious virions. We also tested these drugs on KSHV+ PEL cell line, BCP-1, and found that all of drugs except Tocilizumab displayed some inhibitory effects on BCP-1 cell growth at high concentrations (**Fig. 3A**). By using non-toxic concentrations, we found again that Azithromycin and Nafamostat mesylate significantly increased viral lytic (but not latent) gene expression from BCP-1 cells (**Figs. 3B & S3**). Surprisingly, one more drug, Remdesivir (an adenosine analog) also displayed induced effects on BCP-1 cells, which were not seen on iSLK.219 cells.

**Figure 2.**
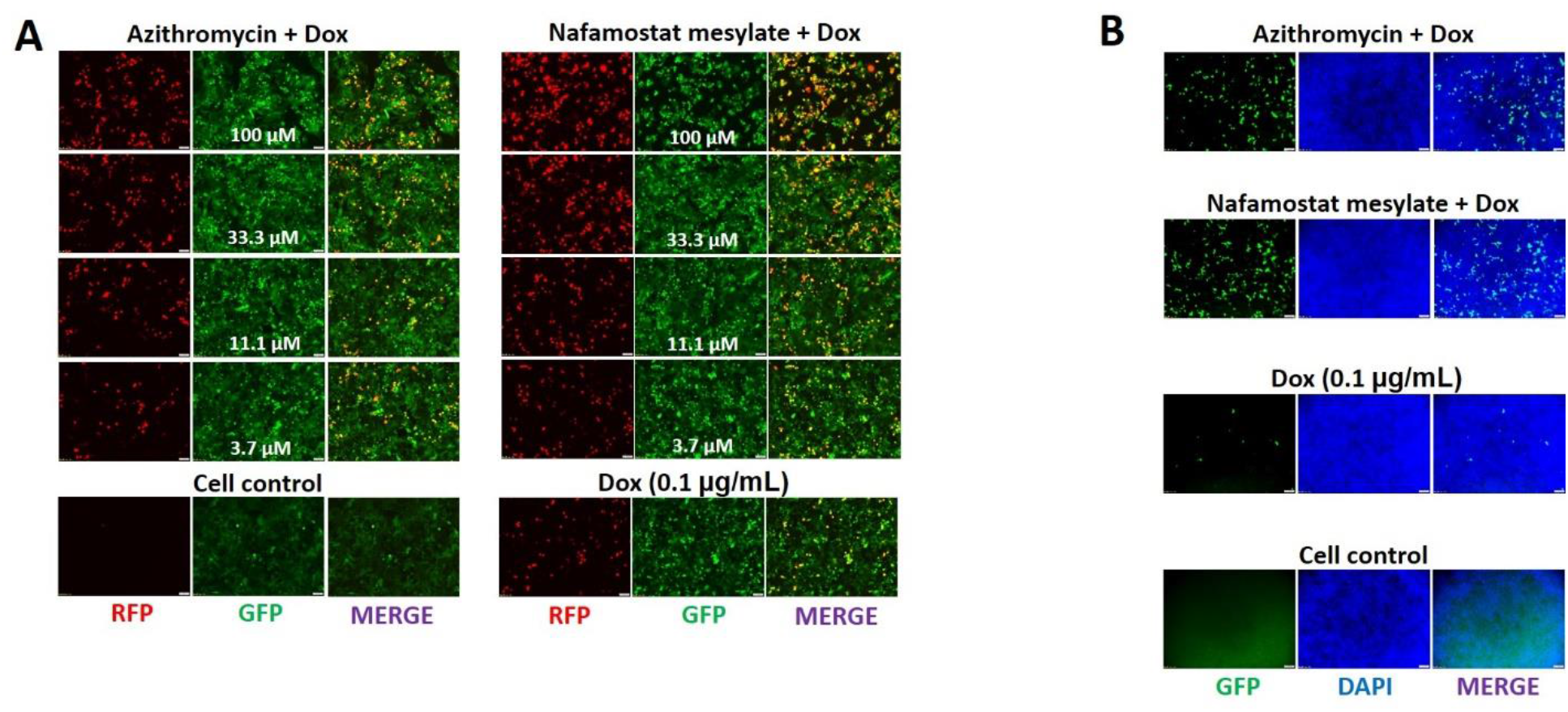
Some drugs currently used for anti-COVID-19 treatment are able to induce KSHV lytic reactivation. (**A**) The iSLK.219 cells were treated with a dose range of anti-COVID-19 drugs, Azithromycin and Nafamostat mesylate, with doxycycline (Dox, 0.1 μg/mL) induction for 72 h. The expression of RFP and GFP were detected using fluorescence microscopy. (**B**) The supernatants from iSLK.219 cells treated by Azithromycin or Nafamostat mesylate for 72 h (11.1 μM, respectively) in combination with Dox were collected to infect naïve HEK293T cells, then GFP expression was detected using fluorescence microscopy.

**Figure 3.**
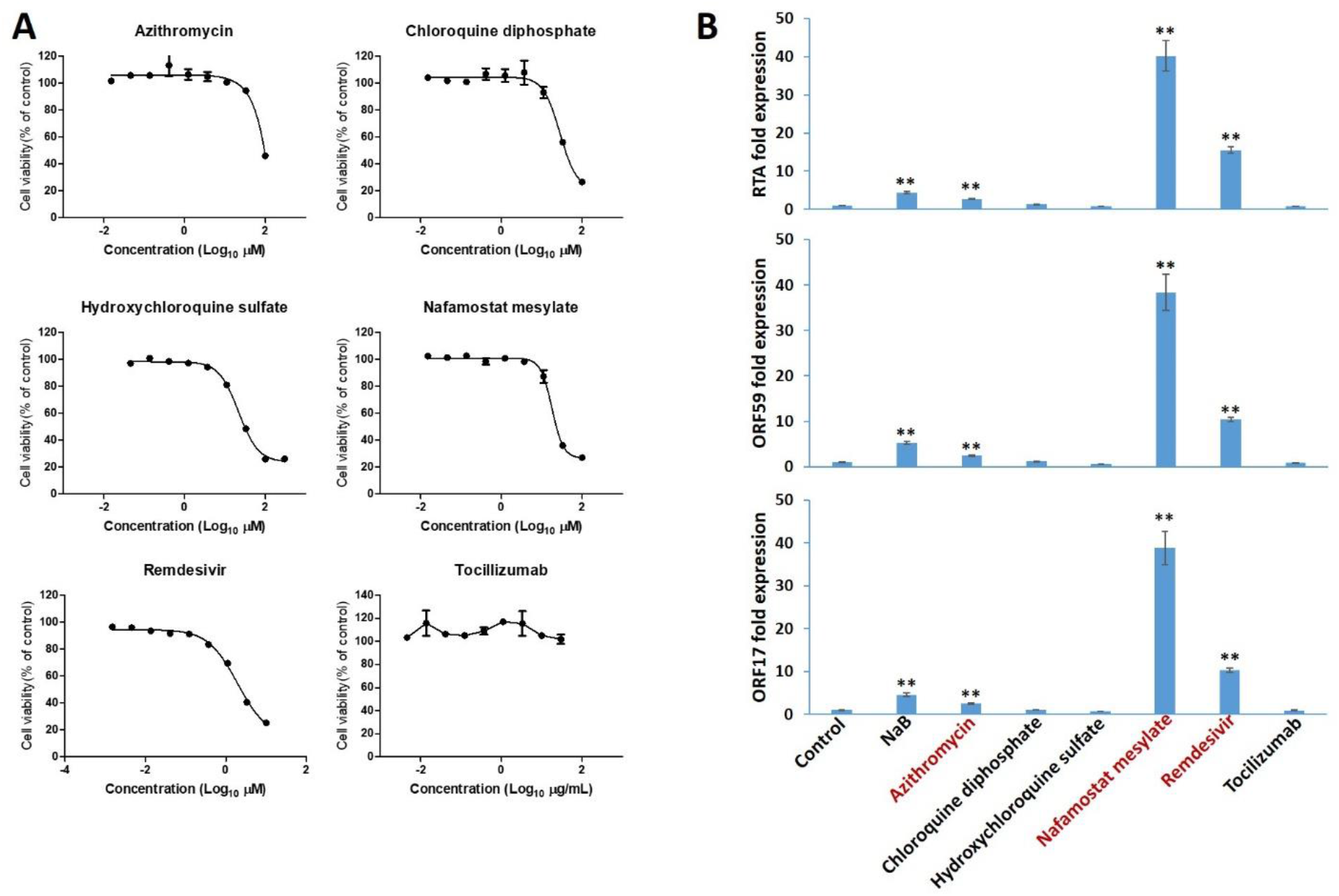
The impacts of anti-COVID-19 drugs on the growth and viral gene expression of KSHV+ tumor cells. (**A**) BCP-1 cells were treated with a dose range of 6 anti-COVID-19 drugs, for 72 h, then the cell proliferation status was examined using the WST-1 cell proliferation assays (Roche). (**B**) BCP-1 cells were treated with Azithromycin (10 μM), Chloroquine diphosphate (10 μM), Hydroxychloroquine sulfate (10 μM), Nafamostat mesylate (10 μM), Remdesivir (3 μM), Tocilizumab (20 μg/mL), respectively, for 72 h, then the transcripts of representative lytic genes were quantified by using qRT-PCR. The sodium butyrate (NaB, 0.3 mM) was used as a positive control. Error bars represent S.D. for 3 independent experiments, ** = p<0.01 (*vs* the vehicle control).

Some major intracellular signaling pathways such as MAPK and NF-κB have been reported to closely regulate KSHV lytic reactivation (Broussard and Damania, 2020b). Here we found that treatment with Azithromycin upregulated MAPK signaling activities (increased phosphorylation of JNK, ERK and p38 kinases), while Nafamostat mesylate treatment downregulated NF-κB activity (reduced phosphorylation of p65) from KSHV+ PEL cells (**Fig. 4A**). To further determine if these signaling pathways are responsible for anti-COVID-19 drugs induced viral lytic reactivation, we first pre-treated iSLK.219 cells with U0126, a specific MAPK kinase inhibitor, which dramatically blocked Azithromycin induced viral lytic reactivation (**Fig. 4B**). Next, the iSLK.219 cells were transfected with a construct expressing NF-κB p65 (pcFLAG-p65 at 0.1 or 0.4 μg, respectively), which effectively blocked Nafamostat mesylate induced viral lytic reactivation in a dose-dependent manner when compared to vector control (**Fig. 4C**). Furthermore, TNF-α, a strong NF-κB inducer treatment was also able to reduce viral lytic reactivation caused by Nafamostat mesylate (**Fig. S4**). Together, these data demonstrate that some anti-COVID-19 drugs can induce KSHV lytic reactivation from latently infected cells through manipulation of intracellular signaling pathways.

**Figure 4.**
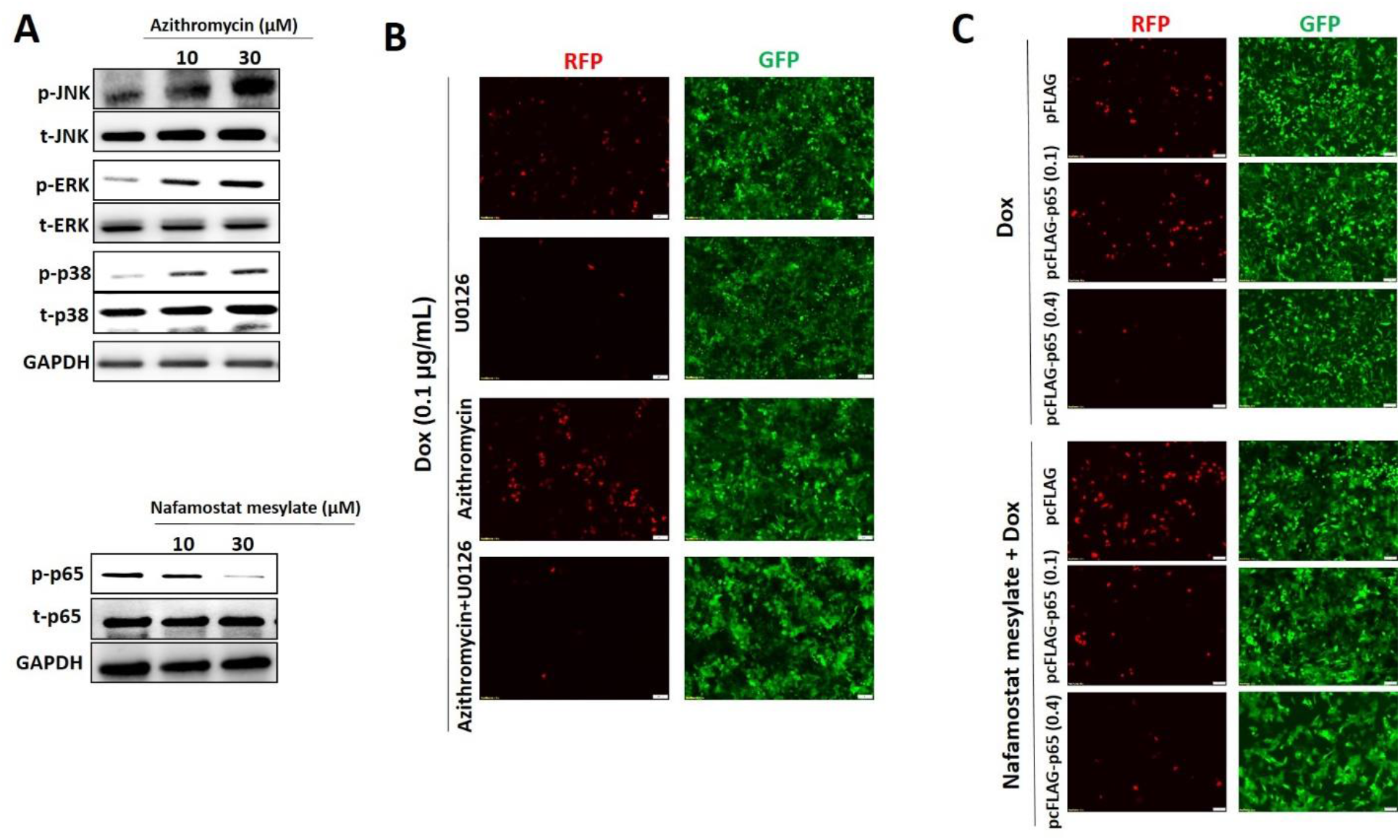
Cellular mechanisms of Azithromycin and Nafamostat mesylate induced KSHV lytic reactivation. (**A**) BCP-1 cells were treated with indicated concentrations of Azithromycin or Nafamostat mesylate for 72 h, then protein expression was measured by immunoblots. Representative blots from one of two independent experiments were shown. (**B**) The iSLK.219 cells were pre-treated with U0126 (10 μg/mL), a MAPK kinase inhibitor, for 12 h, then addition of Azithromycin together with doxycycline (Dox, 0.1 μg/mL) induction for 72 h. (**C**) The iSLK.219 cells were transiently transfected with a construct expressing NF-κB p65 (pcFLAG-p65 at 0.1 or 0.4 μg, respectively) or control vector, then addition of Nafamostat mesylate together with Dox induction for 72 h. The expression of RFP and GFP were detected using fluorescence microscopy.

In the current study, we report for the first time that SARS-CoV-2 encoded proteins and some anti-COVID-19 drugs were able to induce lytic reactivation of KSHV, one of major human oncogenic viruses. These events may facilitate KSHV dissemination as well as initiate viral oncogenesis in those KSHV+ patients exposed to COVID-19 and the related treatments, especially in the case of immunosuppressed patients. Therefore, these patients need follow-ups to monitor KSHV viral loads and virus-associated malignancies development risks, even after they have fully recovered from COVID-19. There are a few limitations and unanswered questions in our study. Firstly, we did investigate using the live virus of SARS-CoV-2 because we do not have access. However, since the S and N proteins represent major and abundant structural proteins of SARS-CoV-2, it will not be surprising to see the live virus display similar effects on this induction. Secondly, we do not have clinical data to support our findings yet. However, this is difficult because the KSHV test is not a routine examination in COVID-19 patients. Further, the seroprevalence of KSHV infection in the general population of the United States is less than 10%, but in most of sub-Saharan Africa, the overall seroprevalence is more than 50% (Mesri et al., 2010). Since mother-to-child transmission of KSHV through saliva is the most common route of transmission, there is a high prevalence of early childhood KSHV infection in some areas (Cao et al., 2014; El-Mallawany et al., 2019; Newton et al., 2018). Moreover, KS has now become one of the most common overall childhood cancers throughout the central, eastern, and southern regions of Africa (El-Mallawany et al., 2018). As we know, children are also susceptible to SARS-CoV-2 infection, although with milder symptoms and lower mortality rates (Bogiatzopoulou et al., 2020). In addition, the seroprevalence of KSHV infection is greatly increased in other sub-populations such as HIV-infected individuals, homosexual men and organ transplant recipients (Mesri et al., 2010). Interestingly, a recent case reported the reactivation of several herpesviruses in a small number of COVID-19 patients (Drago et al., 2020; Xu et al., 2020; Tartari et al., 2020). Thirdly, it remains unclear whether pre-existing KSHV may affect SARS-CoV-2 infection, especially because it is a member of the herpesvirus family which can easily establish lifelong infection in a host. SARS-CoV-2 infects human cells through interaction with the Angiotensin-converting enzyme 2 (ACE2) receptor (Hoffmann et al., 2020), in contrast, there is no data reporting KSHV regulation of ACE2 expression. We previously demonstrated that KSHV *de novo* infection upregulated the expression of one multifunctional glycoprotein, CD147 (also named as Emmprin or Basigin), which was also strongly expressed in KS tissues (Qin et al., 2010; Dai et al., 2015). Interestingly, CD147 was recently found to be one of co-receptors to facilitate SARS-CoV-2 entry and invasion of host cells (Wang et al., 2020). Therefore, it will be interesting to explore the potential association between KSHV and these SARS-CoV-2 receptors or co-receptors in different types of host cells.

## Methods

### Cell culture and reagents

Human iSLK.219 cells are latently infected with a recombinant rKSHV.219 virus and contain a doxycycline (Dox)-inducible RTA, constructed and named by Dr. Don Ganem’s lab (Myoung and Ganem, 2011). The rKSHV.219 virus expresses both the red fluorescent protein (RFP) under the control of KSHV lytic PAN promoter and the green fluorescent protein (GFP) under the control of the elongation factor 1 promoter (EF-1α) (Vieira and O’Hearn, 2004). HEK293T (Human embryonic kidney 293T) cells and KSHV+ PEL cell line, BCP-1, were purchased from American Type Culture Collection (ATCC) and cultured as recommended by the manufacturer. The anti-COVID-19 drugs: Azithromycin, Chloroquine diphosphate, Hydroxychloroquine sulfate, Nafamostat mesylate, Remdesivir, and Tocilizumab were purchased from Selleck Chemicals. All the other chemicals if not indicated specifically were purchased from Sigma-Aldrich.

### Plasmids transfection

Human iSLK.219 cells were transfected with recombinant vectors of SARS-CoV-2 spike protein (S), nucleocapsid protein (N) (both purchased from Sino Biological), pCR3.1-RTA (a gift from Dr. Yan Yuan, University of Pennsylvania) (Chen et al., 2019), pFLAG-CMV2-p65 (pFLAG-p65, a gift from Dr. Ren Sun, University of California Los Angeles) (DeFee et al., 2011), or vector controls, using Lipofectamine™ 3000 reagent (Invitrogen). Transfection efficiency was normalized through co-transfection of a lacZ reporter construct and determination of β-galactosidase activity using a commercial β-galactosidase enzyme assay system (Promega).

### qRT-PCR

Total RNA was isolated using the RNeasy Mini kit (Qiagen), and cDNA was synthesized using a SuperScript III First-Strand Synthesis SuperMix Kit (Invitrogen). Primers used for amplification of target genes are listed in **Table S1**. Amplification was carried out using an iCycler IQ Real-Time PCR Detection System, and cycle threshold (Ct) values were tabulated in duplicate for each gene of interest in each experiment. “No template” (water) controls were used to ensure minimal background contamination. Using mean Ct values tabulated for each gene, and paired Ct values for *β-actin* as a loading control, fold changes for experimental groups relative to assigned controls were calculated using automated iQ5 2.0 software (Bio-rad).

### Cell proliferation assays

Cell proliferation was measured using the WST-1 Assay (Roche). Briefly, after the period of treatment of cells, 10 μL/well of cell proliferation reagent, WST-1 (4-[3-(4-Iodophenyl)-2-(4-nitro-phenyl)-2H-5-tetrazolio]-1,3-benzene disulfonate), was added into 96-well microplate and incubated for 3 h at 37°C in 5% CO_2_. The absorbance of samples was measured using a microplate reader at 490 nm.

### Western blot

Total cell lysates (20 μg) were resolved by 10% SDS–PAGE, transferred to nitrocellulose membranes, and immunoblotted with antibodies to SARS-CoV-2 S or N (Abcam), phosphor (p)-p65/total (t)-p65, p-ERK/t-ERK, p-JNK/t-JNK, p-p38/t-p38 and GAPDH as a loading control (Cell Signaling). Immunoreactive bands were identified using an enhanced chemiluminescence reaction (Perkin-Elmer), and visualized by autoradiography.

### Statistical analysis

Significance for differences between experimental and control groups was determined using the two-tailed Student’s *t*-test.

## Acknowledgements

This work was supported by NIH/NCI R01CA228166. Additional support was provided in part by the Arkansas Bioscience Institute, the major research component of the Arkansas Tobacco Settlement Proceeds Act of 2000. Funding sources had no role in study design, data collection and analysis, decision to publish, or preparation of the manuscript.

## Author contributions

J.C., L.D. and L.B. conducted experiments. J.C. and Z.Q. performed data analysis. Z.Q. designed research. S.R.P. and Z.Q. directed research as well as wrote the manuscript.

## Data Availability

All relevant data are within the manuscript and its supplemental files.

