## Supplemental Table and Figure for "SARS-CoV-2 proteins and anti-COVID-19 drugs induce lytic reactivation of an oncogenic virus"

**Supplemental Table 1. Primer sequences for qRT-PCR.**

| Gene | Sequences (5' → 3') |
|---|---|
| *LANA* | *sense TCCCTCTACACTAAACCCAATA* |
| | *antisense TTGCTAATCTCGTTGTCCC* |
| *RTA* | *sense TAATGTCAGCGTCCACTCC* |
| | *antisense TTCTGGCACGGTCAAAGC* |
| *ORF59* | *sense CGAGTCTTCGCAAAAGGTTC* |
| | *antisense AAGGGACCAACTGGTGTGAG* |
| *ORF17* | *sense AGATTTTTCACGGGGGCTCTGG* |
| | *antisense TGGGCTGGACACTGGGTCTATTTC* |
| *β-actin* | *sense GGAAATCGTGCGTGACATT* |
| | *antisense GACTCGTCATACTCCTGCTTG* |

**Supplemental Figure Legend.**

**Figure S1. Ectopic expression of SARS-CoV-2 proteins in KSHV latently infected cells.** The iSLK.219 cells were transfected with vector control or vectors encoding SARS-CoV-2 spike protein (S), nucleocapsid protein (N) for 72 h, then protein expression was detected using Western blot.

**Figure S2. The impacts of anti-COVID-19 drugs on KSHV lytic reactivation.** The iSLK.219 cells were treated with a dose range of anti-COVID-19 drugs together with doxycycline (Dox, 0.1 μg/mL) induction for 72 h. The expression of RFP and GFP were detected using fluorescence microscopy.

**Figure S3. The impacts of anti-COVID-19 drugs on viral latent gene expression from KSHV+ tumor cells.** BCP-1 cells were treated with Azithromycin (10 μM), Chloroquine diphosphate (10 μM), Hydroxychloroquine sulfate (10 μM), Nafamostat mesylate (10 μM), Remdesivir (3 μM), Tocilizumab (20 μg/mL), respectively, for 72 h, then the transcripts of representative latent gene, *Lana*, were quantified by using qRT-PCR. The sodium butyrate (NaB, 0.3 mM) was used as a positive control. Error bars represent S.D. for 3 independent experiments.

**Figure S4. Pre-treatment of TNF-α blocks Nafamostat mesylate induced KSHV lytic reactivation.** The iSLK.219 cells were pre-treated with TNF-α (0.2 μg/mL) for 12 h, then addition of Nafamostat mesylate together with doxycycline (Dox, 0.1 μg/mL) induction for 72 h. The expression of RFP and GFP were detected using fluorescence microscopy.

**Figure S1**

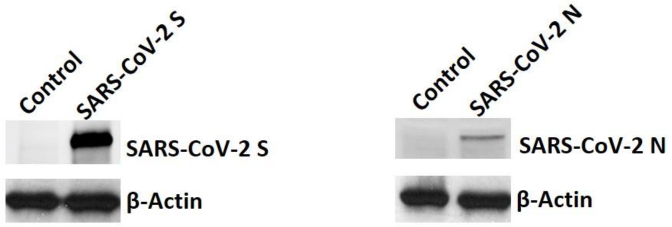
Control
SARS-CoV-2 S
SARS-CoV-2 S
β-Actin
Control
SARS-CoV-2 N
SARS-CoV-2 N
β-Actin

### Figure S2

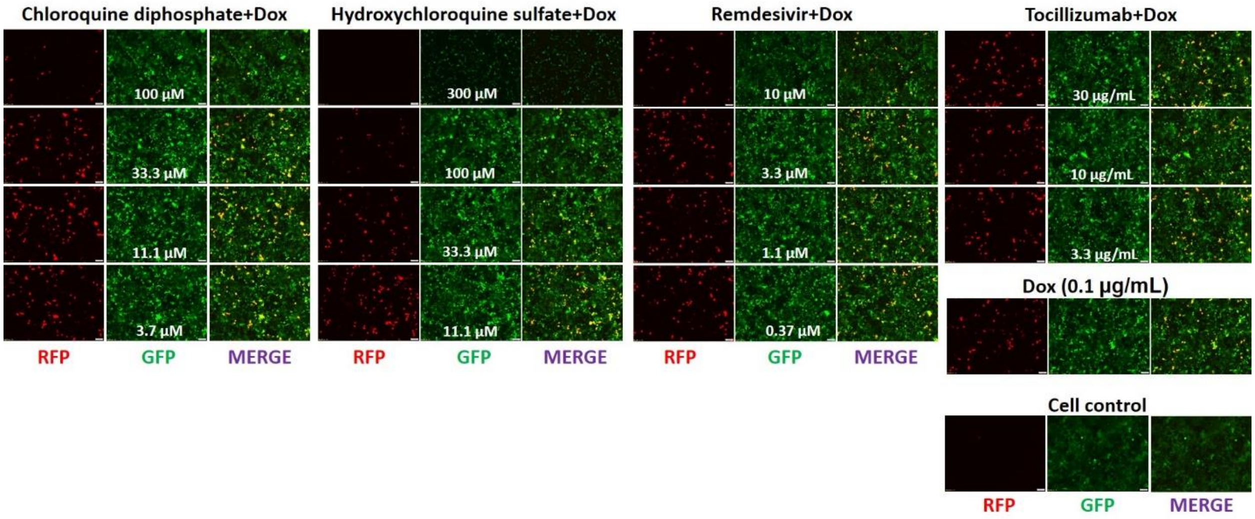
Chloroquine diphosphate+Dox
100 μM
33.3 μM
11.1 μM
3.7 μM
Hydroxychloroquine sulfate+Dox
300 μM
100 μM
33.3 μM
11.1 μM
Remdesivir+Dox
10 μM
3.3 μM
1.1 μM
0.37 μM
Tocillizumab+Dox
30 μg/mL
10 μg/mL
3.3 μg/mL
Dox (0.1 μg/mL)
Cell control
RFP
GFP
MERGE

**Figure S3**

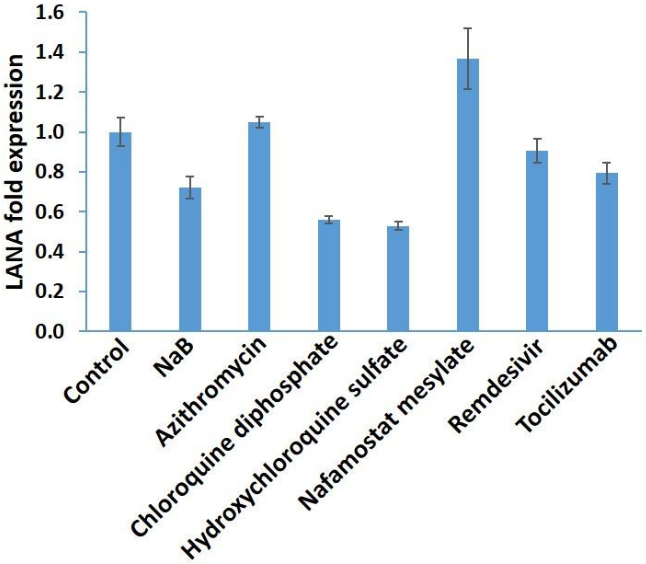
LANA fold expression
0.0
0.2
0.4
0.6
0.8
1.0
1.2
1.4
1.6
Control
NaB
Azithromycin
Chloroquine diphosphate
Hydroxychloroquine sulfate
Nafamostat mesylate
Remdesivir
Tocilizumab

**Figure S4**

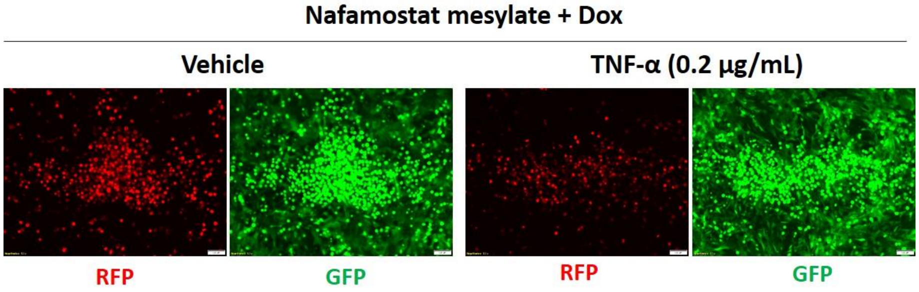

Nafamostat mesylate + Dox
Vehicle
TNF-α (0.2 μg/mL)
RFP
GFP
RFP
GFP
